# Impaired dynamic profile of brain recurring functional connectivity states in Alzheimer’s disease

**DOI:** 10.1101/2020.10.03.324673

**Authors:** Somayeh Maleki-Balajoo, Davud Asemani, Ali Khadem, Hamid Soltanian-Zadeh

## Abstract

Up to now, disrupted functional organization of the brain in Alzheimer’s disease (AD) has mostly been examined by assessing “static” functional connectivity (FC). However, recent studies have provided clear evidence that brain FC exhibits intrinsic spatiotemporal “dynamic” organization also being significantly affected in AD. However, inconsistency in former studies is a motivation for further investigation of impaired dynamic profile of brain FC networks in Alzheimer’s disease._In this work, we identified recurring FC states during rest (eyes-open) in 24 AD patients and 37 healthy controls (HC) by combining sliding window-based method (examining dynamic FC) and k-means clustering algorithm (identifying recurring FC states over time). To define the optimum number of recurring FC states in k-means algorithm, we employed non-supervised validity criteria such as silhouette value and Dunn index over bootstrap samples. Then, we examined differences in dynamic properties of recurring FC states including probability of occurrence, lifetime and switching profile between groups. Afterward, association between impaired dynamic profile of recurring FC states and cognitive performance was assessed in AD patients. Our findings revealed three recurring FC states and among them, FC state with the most significant differences in probability of occurrence (corrected p-value < 0.0001) included brain regions involved in default mode, frontoparietal and salience networks. This recurring FC state occurred significantly less and lasted shorter in patients with AD than in the HC. Further, the probability of occurrence in this recurring FC state significantly correlated positively with cognitive decline in patients with AD. Our findings suggest more decreased in cognitive performance in patients with AD associated with more reduced ability of AD patients to access clinically relevant impaired brain FC networks. Impaired dynamic profile of recurring FC states in AD provide greater understanding of AD’s pathophysiology.

## 1- Introduction

Alzheimer’s disease (AD) is the most prevalent neurodegenerative disorder, which has become a major public health problem in the recent decades (Prince et al., 2016). Recent studies have exploited resting-state functional magnetic resonance imaging (rs-fMRI) to investigate the functional connectivity (FC) impairments in AD. Disruption of several resting-state networks in AD have been reported, including the default mode network (DMN), involved in memory formation and retrieval (Andrews-Hanna, 2012) and self-related processing (L. Zhang et al., 2020), frontoparietal network (FPN), flexibly controlling and interacting with other brain networks (Marek & Dosenbach, 2018), the salience network (SN), directing attention toward or away from internal processing in concert with the DMN (Menon & Uddin, 2010), somatomotor network (SMN) and visual network (Mosimann, Felblinger, Ballinari, Hess, & Muri, 2004). Previous studies have ignored dynamic nature of FC and mostly focused on assessing ‘static’ FC which represents mean connectivity over the period of scanning (Damoiseaux, Prater, Miller, & Greicius, 2012; Greicius, Srivastava, Reiss, & Menon, 2004; Sorg, Riedl, Perneczky, Kurz, & Wohlschlager, 2009; Zhao, Lu, Metmer, Li, & Lu, 2018). Most recently, several studies have provided empirical evidences in healthy subjects (Cabral et al., 2017; Larabi et al., 2020; Maleki Balajoo, Asemani, Khadem, & Soltanian-Zadeh, 2020) and psychiatric (Figueroa et al., 2019; Sakoglu et al., 2010) and neurological (Fu et al., 2019; Gu et al., 2020; Jones et al., 2012; Kim et al., 2017; Niu et al., 2019; Schumacher et al., 2019; Sourty et al., 2016) disorders that not only the intrinsic brain FC at rest is dynamic during the period of scanning (Allen et al., 2014; Betzel, Fukushima, He, Zuo, & Sporns, 2016; Chang & Glover, 2010; Hutchison et al., 2013), but also temporal properties of FC reconfiguration over time are associated with symptoms, behavioral and cognitive performance (Cabral et al., 2017; Figueroa et al., 2019; Gu et al., 2020; Larabi et al., 2020; Tian, Li, Wang, & Yu, 2018; Viviano, Raz, Yuan, & Damoiseaux, 2017). For AD, few studies have investigated dynamic functional connectivity (dFC) impairments or reconfiguration of brain FC states over time by using rs-fMRI data (Fu et al., 2019; Gu et al., 2020; Jones et al., 2012; Schumacher et al., 2019). Among those studies, Jones et al. (2012) hypothesized that variability in interstice FC networks is related to non-stationary nature of FC within interstice brain FC networks and mainly investigated the dynamic property of brain’s modular organization while focusing only on DMN in AD for validating their hypothesis (Jones et al., 2012). Findings from other studies suggest temporal properties of recurring FC states are generally disrupted in AD, but detailed findings are not convergent. For example, Gu et al., (2020) and Fu et al., (2019) reported patients with AD spent less time in baseline (global) state compared with HC subjects, while Schumacher et al (2019) reported an opposite result using a similar method proposed in (Fu et al., 2019). On the other hand, Gu et al., (2020) reported significant association between impaired dFC and cognitive performances in patients with AD, while Schumacher et al., (2019) indicated no significant relation between impaired dFC and clinical symptom severity. Employing different approaches to estimate dFC and using different datasets may have resulted in divergent findings. The inconsistency between results of previous studies necessitates further investigation of dFC in AD.

In the current study, we used a recently developed method based on sliding-window analysis, in which an exponentially tapered function proposed by (Pozzi, Di Matteo, & Aste, 2012) and used in (Betzel et al., 2016; Zalesky, Fornito, Cocchi, Gollo, & Breakspear, 2014) to estimate dFC pattern in each window by calculating the weighted Pearson product-moment correlation between brain regions. Sliding-window analysis is the most commonly used method to calculate consecutive dFC matrices over time (Hutchison et al., 2013; Sakoglu et al., 2010) because of both easy implementation and simple interpretation. Here, we employed dFC analysis to investigate whether there are specific configurations of FC states over time that can differentiate patients with AD from HC subjects.

In this research, we first summarized each dFC matrix by extracting its leading eigenvector. Then, we utilized the k-means algorithm on concatenated leading eigenvectors extracted from all subjects in both healthy control (HC) and AD patient groups to identify brain recurring FC states that reoccur over time. We employed non-supervised validity criteria such as silhouette value and Dunn index over bootstrap samples to define the optimum number of FC states in k-means algorithm. Afterward, statistical group comparisons on the dynamic properties of the identified brain recurring FC states at the optimum cluster solution including probability of occurrence, lifetime and switching profiles to explore how AD effects the dynamic profile of brain recurring FC states. Then, we defined the most significant recurring FC state as the FC state with the strongest group difference in probability of occurrence with smallest p-value. In the following to explore that identified most significant recurring FC state in the optimum level of clustering is independent of the level of clustering, we identified the most significant recurring FC state in all level of clustering. The number of clusters varies from 2 to 17. Then, we evaluated the association between dynamic characteristic of identified recurring FC states in AD and cognitive performances in patients with AD. We hypothesized that alterations in dynamic profile of identified recurring FC states in AD patients would be positively correlated with cognitive impairment in AD (Buckner, Andrews-Hanna, & Schacter, 2008; Filippi, Spinelli, Cividini, & Agosta, 2019; Pasquini et al., 2019; Sorg et al., 2009).

## 2- Materials and Methods

### 2-1- Participants

The imaging data was obtained from the Alzheimer’s Disease Neuroimaging Initiative (ADNI) (http://adni.loni.usc.edu)) representing one of the best openly accessible Magnetic Resonance Imaging (MRI) datasets of dementia and healthy control (HC). The sample used in this study consisted of HC subjects (n = 46) and patients with Alzheimer’s disease (AD) (n = 25) which all completed both structural MRI and resting-state functional MRI (rs-fMRI) scans. Subjects should have eyes open during rs-fMRI scans. Visually inspection of all the structural MRI and rs-fMRI data for imaging artifacts and excessive head motions, identified seven HC subjects which were excluded from this study. A set of 64 subjects (nHC = 39 and nAD = 25) with high-quality images were used for further analysis in this study. HC subjects were in a preclinical stage in which they showed no signs of depression, significant memory concern, mild cognitive impairment, or dementia (see demographic information in Table 1).

**Table 1-.** Demographic information.

| Variables | HC | AD | p-value |
| --- | --- | --- | --- |
| No. of subjects | 37 | 24 | - |
| Mean age in years ± SD | 75.3 ± 6.3 | 72.26 ± 7.65 | 0.096 |
| Sex (M/F) | 12/25 | 7/17 | 0.988 |
| Mean years of education ± SD | 16.59 ± 1.97 | 15.79 ± 2.79 | 0.194 |
| Handedness (R/L) | 36/1 | 22/2 | 0.698 |
| Mean MMSE* score ± SD | 28.92 ± 1.42 | 21.87 ± 2.85 | <0.001** |

Comparing Hypothesis test: for variables including age, years of education and MMSE score, we used two-sample t-test to compare means. Ut in case of binary variables such as sex and handedness, Chi-square test was used to compare proportions. AD = Alzheimer's disease; HC = Healthy control subjects; SD = standard deviation. \*: Mini-Mental State Examination; \*\* p < 0.05.

### 2-2- Data acquisition protocols

MRI scanning of all subjects included in this study was performed by Philips Medical System scanners with a 3T field strength at baseline of ADNI-2 phase according to ADNI protocol (Jack et al., 2008) to reduce the effect of using different MR scanners. The T1-weighted structural images were acquired using magnetization-prepared rapid-acquisition gradient echo (MP-RAGE) pulse sequence with the following parameters: 170 sagittal slices (gap 1.2 mm); TR=6.8 milliseconds (msec); TE=3.1 msec; flip angle=9°; acquisition matrix = 256×256; voxel size = 1.2×1.0×1.0 mm3. The rs-fMRI data (eye-open) consist of 140 volume images acquired using a gradient-echo Echo planar imaging (EPI) pulse sequence: 48 axial slices (gap 3.3 mm); TR= 3000 msec; TE=30 msec; flip angle=80°; acquisition matrix = 64×64; voxel size = 3.31×3.31×3.31 mm3.

### 2-3- Data preprocessing

The rs-fMRI data were preprocessed using Statistical Parametric Mapping (SPM12) and the Data Processing Assistant for Resting-State fMRI toolbox (DPARSF) (Yan, Wang, Zuo, & Zang, 2016). After converting the DICOM images into 3D-NIFTI, the first three volumes of each subject’s EPI images were removed as the BOLD signal transient state, and the remaining volumes were rigidly registered to the same subject’s mean EPI-image using affine transformation (with 6 degrees of freedom) to compensate for the head motion. Three subjects, one patient with AD and two HC subjects, were excluded due to excessive movement (cumulative translation >2 mm or rotation >2°), so a set of 61 subjects (n_HC_ = 37 and n_AD_ = 24) were approved for final analysis in this study. Then, T1-weighted images were segmented based on tissue-probability maps into gray matter (GM), white matter (WM) and cerebrospinal fluid (CSF) using the ‘mixture model’ clustering algorithm (Y. Zhang, Brady, & Smith, 2001). Afterward, rs-fMRI data were registered onto the Montreal Neurological Institute (MNI) template using the SPM12 unified segmentation on the T1 image routine. MNI tissue-probability maps of GM, WM, and CSF were warped onto single-subject T1-weighted images and were stored as masks for later use during the nuisance covariate regression. All of the EPI images were resampled to an isotropic voxel size of 3×3×3 mm^3^ and spatially smoothed by using a Gaussian filter (FWHM Gaussian kernel 4× 4× 4 mm^3^). Then, linear trends were removed and the rs-fMRI data were temporally filtered with a band-pass filter (0.01-0.1 Hz) to reduce low and high frequency noises and extract the physiologically desired signals (Lu et al., 2007). Finally, the nuisance variables including motion parameters with the Friston 24-parameter model, WM, and the CSF signals (Friston, Williams, Howard, Frackowiak, & Turner, 1996; Kelly, Uddin, Biswal, Castellanos, & Milham, 2008) were also regressed out from the rs-fMRI data to eliminate their effects on the BOLD fluctuations.

### 2-4- Dynamic functional connectivity

The regional time series were extracted from all 112 brain regions which were parcellated based on Harvard-Oxford atlas (Bohland, Bokil, Allen, & Mitra, 2009) (Figure 1A). we computed dFC between each pair of regional time series using a tapered sliding-window approach (Zalesky et al., 2014) (Figure 1B). To do so, the regional time series were firstly divided into overlapping windows of approximately 100 seconds (sec) and step of one TR (3 sec). With a sampling rate of TR=3 sec, the windows were represented by 33 samples (window length = 99 sec) and 32 samples overlap between successive windows. The reason behind the selection of a window of approximately 100 sec length was that it is the shortest possible window to capture a full cycle of the slowest frequency component due to high-pass cutoff frequency (0.01 Hz) (Betzel et al., 2016; Leonardi & Van De Ville, 2015; Maleki Balajoo et al., 2020; Zalesky & Breakspear, 2015). It is noteworthy that, Gu et al., (2020) utilized a short window length (i.e., 30 sec) to estimate dFC patterns compared with current study, however, window length between 30 and 100 sec has been proposed as a good window length for dFC analysis (Vergara, Abrol, & Calhoun, 2019; Wilson et al., 2015). Then, we defined an exponentially tapered function as proposed by (Pozzi et al., 2012) and used in (Betzel et al., 2016; Zalesky et al., 2014) for each window and calculated the weighted Pearson product-moment correlation between regions *i* and *j* using the observations only within that window. Subsequently, for each subject, the time-resolved dFC matrices of size *N×N×T* were yielded, where *N* = 112 is the number of brain regions and *T = 105* is the number of windows. Finally, at each window of the index *t* (*t* = 1: *T*), the calculated dFC matrix, dFC(t), were z-transformed and vectorized (Figure 1B). In favor of focusing on the fluctuations of FC over time, we removed mean pair-wise correlations over time (Grandjean et al., 2017; Leonardi et al., 2013).

**Figure 1-.**
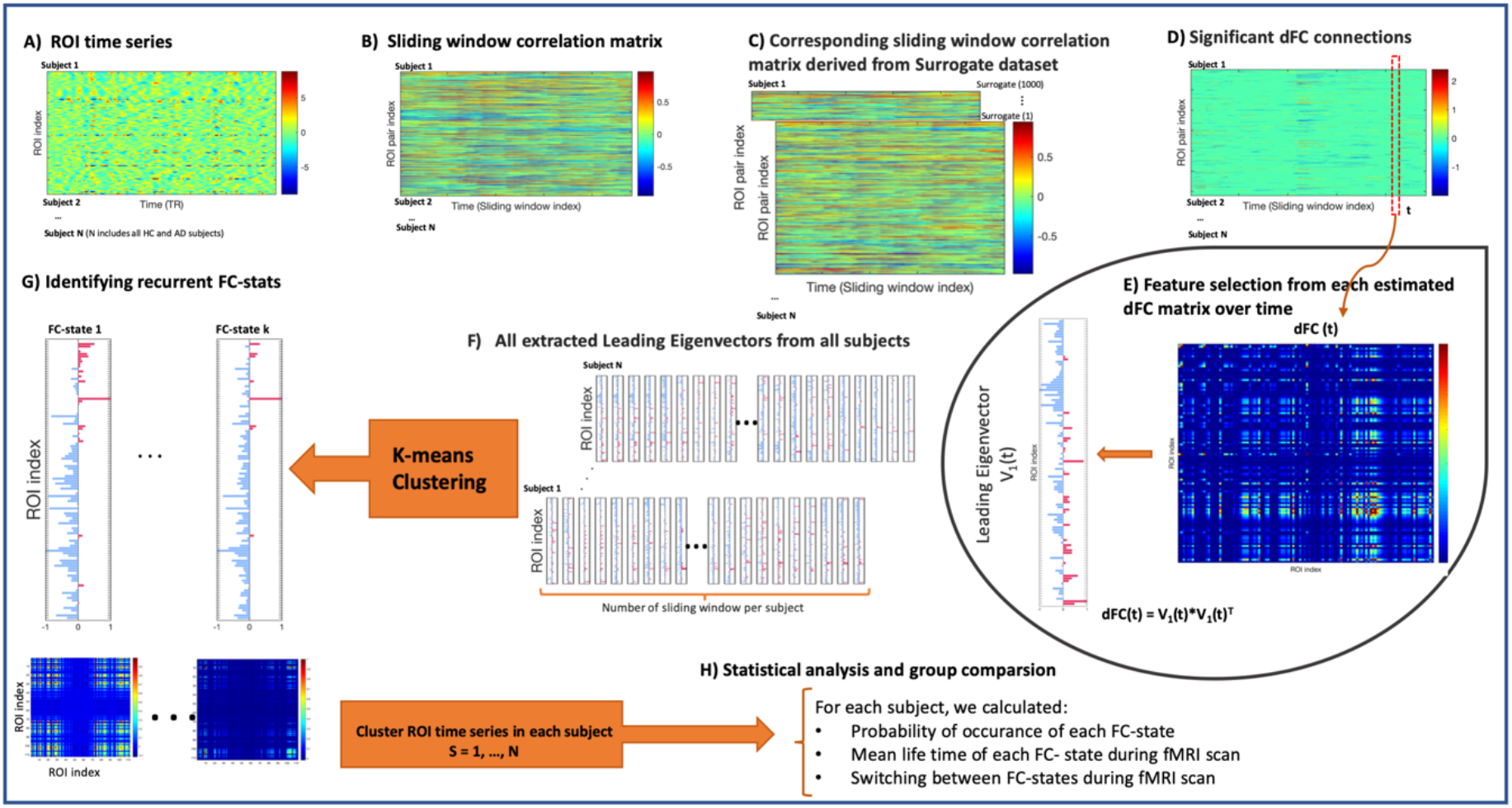
Method summary. (A) For each subject, the time series describing the BOLD signal activity of the Harvard-oxford atlas regions of interest (ROIs) were extracted. (B) The time series were analyzed with sliding-window based approach. Sliding-window correlation matrices describing the degree of whole-brain regional interactions at each time point for each subject are obtained. (C) For each subject, the corresponding sliding-window correlation matrices based on surrogate dataset present comparable features as individual sliding window correlation matrix (B). (D) For each subject, dynamic correlation fluctuations in the surrogate datasets were used to estimate 2.5^th^ and 97.5^th^ confidence intervals for every individual and interactions between ROI pairs. Dynamic correlations outside the confidence intervals are indicated as significant dynamic connections. (E) For each subject, dFC(t) matrix at each time can be defined by column vector in (D) and that would be a N×N matrix, where N is 112 (the number of ROIs in the whole brain). The leading eigenvector V1(t) of each dFC(t) matrix was extracted which is a N×1 vector that explains the most variances in the dFC(t) matrix. If V1(t) is multiplied by its transpose, it reveals the dominant pattern of the dFC(t) matrix. The leading eigenvectors V1 are obtained for each time in all subjects in both groups (HC and AD) and concatenated together, resulting in a large sample leading eigenvector. (F) All extracted leading eigenvectors from all subjects in both groups were clustered into a reduced number of k clusters (here we varied k between 2 and 17). (G) Each cluster is represented by a central vector, which may not necessarily be a member of the data set. We take these cluster centroid vectors as representing FC-state. (H) For each subject, we calculated the probability of occurrence, mean lifetime, and the switching profile of each FC state based on FC state time series. FC-state time series were obtained by defining at each time point, which FC state is the most similar to V_1_(t).

### 2-5- Detecting statistically dynamic connections

Previous studies have shown that dFC estimation approaches and specially sliding window correlation analysis, tend to induce spurious correlations (Hindriks et al., 2016; Kudela, Harezlak, & Lindquist, 2017; Leonardi & Van De Ville, 2015; Maleki Balajoo et al., 2020; Zalesky & Breakspear, 2015). So, for each pair of regions, we performed a statistical hypothesis test to measure the extent of dynamic fluctuations in the calculated correlation coefficients. We generated, for each subject, 1000 phase randomized surrogate time series per ROI such that static FC between every ROI pair was preserved within any set of surrogates, similar to Betzel et al. (2016). For each realization of surrogate data, dFC was estimated by using the tapered sliding-window approach and removing mean pair-wise correlations over time similar to what was applied to the real rs-fMRI time series (Figure 1C). Finally, for each subject and for each dynamic functional connection between every ROI pair, we approximated the null distribution based on all the estimated dynamic fluctuations between that ROI pair from 1000 surrogate data. Then, we assigned the percentile of each dynamic functional connection calculated in each real data based on the corresponding obtained null distribution of dynamic functional connection. A dynamic functional connection was identified as significant dynamic fluctuation with p-value<0.05 if its assigned percentile was less than 2.5^th^ or more than 97.5^th^ percentiles (Figure 1D).

### 2-6- Identifying recurring functional connectivity states

To identify the difference of dFC patterns over time between groups, we first extracted recurrent FC patterns (recurring FC states) in the estimated dFC over time by clustering all the N× N dFC(t) matrices obtained at all windows (time points) across all individuals in both groups. Since dFC(t) matrices are symmetric, each pattern of dFC(t) can be summarized by the upper triangular elements of its matrix including N(N-1)/2 elements (Betzel et al., 2016; Demertzi et al., 2019; Grandjean et al., 2017; Hindriks et al., 2016; Preti, Bolton, & Van De Ville, 2017) or by its leading eigenvector including N elements (Cabral et al., 2017; Figueroa et al., 2019; Larabi et al., 2020). In case of using leading eigenvector, the dimensionality is potentially reduced from N^2^× 1 to N×1 while still explaining most of variance in dFC(t).

In line with previous studies, we used leading eigenvector as a feature reduction approach which captures dominant pattern in each dFC(t) matrix (Cabral et al., 2017; Figueroa et al., 2019; Larabi et al., 2020) (Figure 1E). The leading eigenvector; V_1_, of a symmetric matrix was first introduced and used in community detection in networks (Newman, 2006) to represent the dominant connectivity pattern between nodes of a network and identifies the community structure in the network by clustering all eigenvector elements based on their sign (positive versus negative). The sign was used to label two different types of connections between the communities. The original symmetric matrix can be reconstructed using the outer product as V_1_× V_1_^T^. In the following, we concatenated all the extracted leading vectors, V_1_ (one per window) in all subjects, resulting in a large sample of 6,405 leading eigenvectors ((n_AD_ + n_HC_) * T = (24 + 37) * 105 = 6405 leading vectors) (Figure 1F). . Then, we used k-means clustering approach to partition this large sample into a reduced number of K clusters. Here, we examined 16 levels of clusteringranging from k = 2 to 17 and in each level of clustering, k, the k-means clustering approach divided all samples into k clusters. Each cluster is represented by a central vector (centroid), which may not necessarily be a member of the sample set. These cluster centroids are representative of recurring FC patterns over time which named as recurring FC states (Figure 1G). To obtain the FC pattern of each recurring FC state (cluster), we only need to calculate the outer product of cluster centroid vector and itself. In k-means algorithm, the repetition number was set to 500, which almost doubled the recommended number of 256 repetitions (Nanetti, Cerliani, Gazzola, Renken, & Keysers, 2009), and the iteration number was set to 255 to avoid getting stuck in local optimum, with random initialization in each iteration. The decision to select the range from 2 to 17 for k is that previous works have reported the number of resting-state networks between 7 and 17, depending on the selected criteria (Damoiseaux et al., 2006; Yeo et al., 2011). In the following section, we explain how the optimum level of clustering was determined.

### 2-7- Measurement of Stability and Consistency of clustering to define number of recurring FC states

In this work, we defined the optimum number of FC-states (number of clusters in the clustering approach) based on stability and consistency of the different levels of clustering by measuring internal evaluation criteria in clustering such as the adjusted rand index (ARI), silhouette value and Dunn index over bootstrap samples. We generated 200 bootstrap samples for each group by utilizing bootstrap resampling with replacement approach (Bellec, Rosa-Neto, Lyttelton, Benali, & Evans, 2010) applied to observed dataset and of equal size to the observed data set. For example, in HC group (37 subjects) we created 200 random samples with size of 37. As bootstrapping is resampling with replacement, different versions of the original sample were generated. Afterward, for each of bootstrap samples (we consider one bootstrap sample from each group), we concatenated all the extracted leading vectors of all subjects in HC and AD bootstrapping samples and applied k-means clustering approach to identify recurring FC states while 16 levels of clustering ranging from k = 2 to 17 were examined for each of bootstrap samples. Then, we examined different criteria including adjusted rand index (ARI) (Rand, 1971), silhouette value (Rousseeuw, 1987) and Dunn index (Dunn, 1973) to define the optimum solution. Thus, the optimum number of clusters (K) over all 200 bootstrap samples is selected if at solution K, the higher value for all metrics compared to the previous K-1 solution is acheived or at least the metric values are at least not significantly decreased compared to the previous K-1 solution. The point is that different criteria are not maximized simultaneously and it is usually worthwhile to take a closer look at the performance of the cluster solutions with regard to the different criteria. If you can support a particular cluster solution with one of each of these criteria, this is considered a very strong argument for selecting that cluster solution. In this work, we focused on bootstrapping approach to ensure stability and consistency of recurring FC states across subjects. An ARI value of one indicates that the clustering results across bootstrap samples are identical and a value of zero suggests that clustering results are not similar to each other, whereas negative values indicate a dissimilarity of clustering results higher than chance level. The silhouette value is a measure of how similar a V_1_ vector is to other V_1_ vectors in its own cluster compared with V_1_ vectors in other clusters. This value ranges from -1 to +1. A higher Dunn index indicates better clustering reflected by compact clusters (i.e. small variance within cluster) which are separated well from other near clusters.

### 2-8- Between group comparisons

To assess how the identified recurring FC states during rest varied between patients with AD and HC subjects, we first calculated the probability of occurrence of each recurring FC state (fraction of estimated dFC matrices (time windows) assigned to that recurring FC state throughout the scan duration), the mean lifetime of each recurring FC state (mean number of contiguous estimated dFC matrices (time windows) of the same FC state), and the switching profiles (probabilities of switching from a given FC state to each other one), for each subject. For example, we computed the probability of occurrence of each recurring FC state for each subject as the total number of V1 vectors over time assigned to that given FC state divided by total number of estimated V1 vectors over time for each subject. All values were compared between HC and patients with AD using (nonparametric) permutation-based t-tests (10,000 permutations). For each identified recurring FC state of k FC states obtained by k-means clustering, k hypotheses were tested. To correct for multiple comparison, we adjusted the significance threshold to 0.05/k for Bonferroni correction (green dashed line in Figure 5). In this work, a recurring FC state in optimum cluster solution with the most significant differences in probability of occurrence that yielded the strongest group difference with smallest p-value named as “most significant recurring FC state”. In the following, we explain how we explored the replicability of identified most significant recurring FC state in optimum level of clustering which was significantly differed between patients with AD and controls over levels of clustering (k = 2 to 17).

**Figure 5-.**
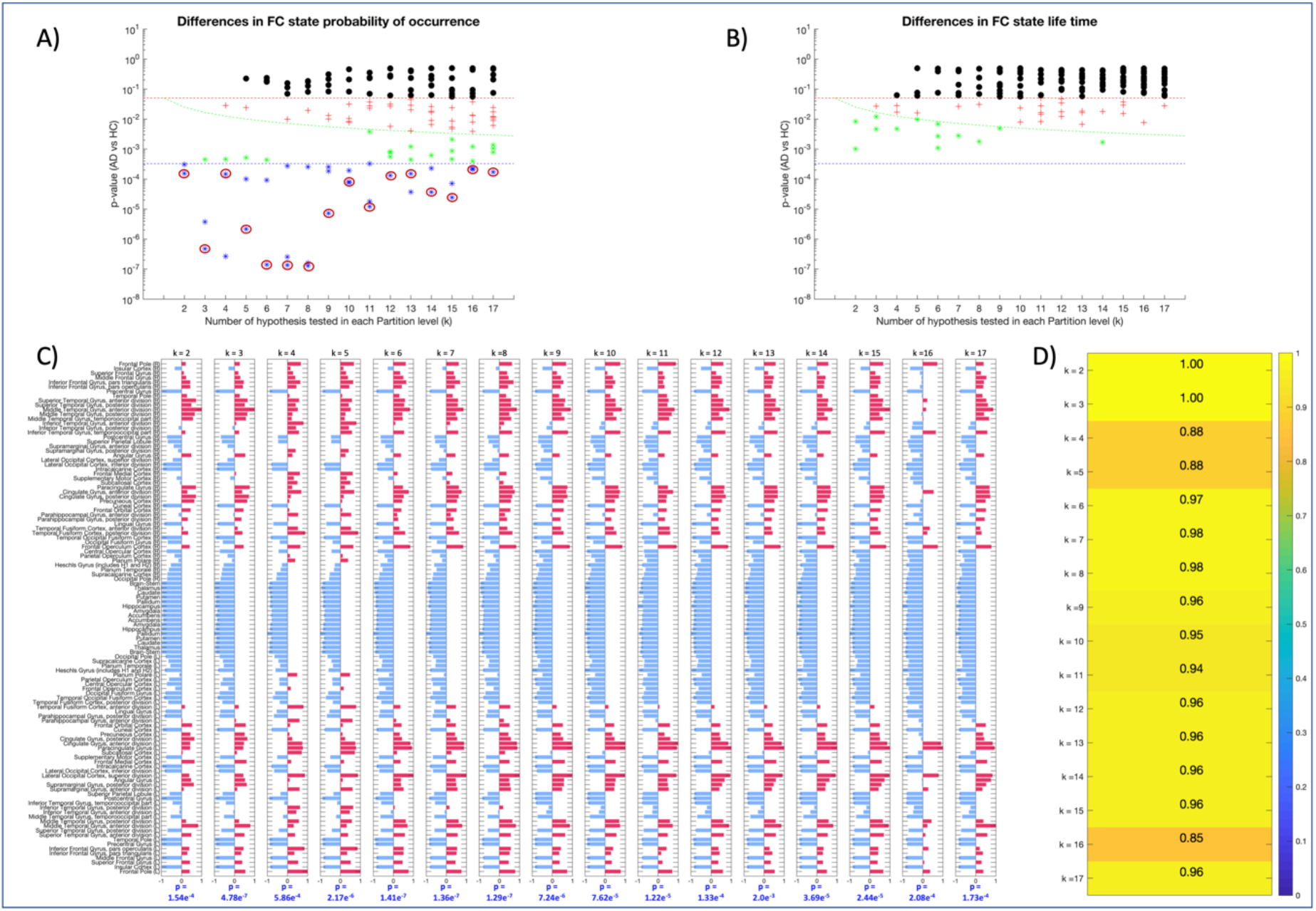
Replicability of FPN-SN-DMN state where the most significant differences in probability of occurrence between patients with AD and healthy control subjects observed, in all levels of clustering. we first computed between-group comparisons in terms of probability of occurrence (A) and lifetime (B) of each FC state for each clustering level of clustering(k = 2:17). We used black dots for cases where there was no significant difference between groups (p-value > 0.05). The most significant results while corrected for all hypothesis in this study by using FWE rate correction 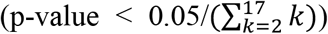 were marked with blue star. The significant results did not survive the correction for all hypothesis while survived the correction for each level of clustering by using FWE rate correction 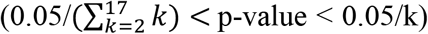 were marked with green star. Finally, the results which p-value is less than 0.05 but do not survive the correction for level of clustering by using FWE rate corrections within each partition model (0.05/k < p-value < 0.05) were marked as red crosses. (C) The circled FC states in the all levels of clustering with red in (A) that occurred significantly less in patients with AD compared to HC consistently represented FPN-SN-DMN networks. (D) correlation between identified FC states in (C) with FPN-SN-DMN state identified in the most stable level of clustering (k =3). FC: functional connectivity; AD: Alzheimer’s disease; FPN: frontoparietal network; SN: salience network; DMN: default mode network; FWE: Family wise error.

### 2-9- Replicability of the most significant recurring FC state

To assess the independency of the identified most significant recurring FC state in the probability of occurrence (in the optimal clustering solution) from what the number of clusters to be, we explored all the identified recurring FC states at all different levels of clustering (k=2:17). Definitely, we assumed that multiple clusters at different levels of clustering might be all meaningful, but at higher levels of clustering, smaller-more grained and rare FC patterns are revealed. In other words, we explored the replicability of the identified most significant recurring FC states in the probability of occurrence in optimum level of clustering over all clustering levels. To do so,

we first computed between-group comparisons in terms of probability of occurrence of each recurring FC state for each clustering level of clustering (k = 2:17). The most significant recurring FC state in each level of clustering marked as a recurring FC state with strongest difference between groups with smallest p-value. Finally, we examined how much the marked FC states in the all levels of clustering correlated with the most significant recurring FC state in the optimal level of clustering.

### 2-10- Correlation analysis for dynamic characteristics of recurring FC states and cognitive decline score

To analyse the relationship of cognitive decline score with dynamic characteristics of recurring FC states, we calculated the Pearson correlation coefficient between mini-mental state examination (MMSE) and dynamic profile of recurring FC state including probability of occurrence and lifetime of recurring FC states and the switching profiles. MMSE score give an overall assessment of subject’s mental status by evaluating memory, the ability of solving simple problems and some thinking skills. The maximum MMSE score is 30 points which achieved mostly by subjects with healthy mental condition. **A score** of **25** or **lower** is considered significant cognitive impairment. On average, the MMSE score of a person with AD declines about two to four points each year (Sheehan, 2012).

## 3- Results

### 3-1- Identifying the most stable level of clustering

We measured stability and consistency of all levels of clustering (k = 2:17) across bootstrap samples with clustering the concatenated V1(t) vectors over time and all subjects (including both HC and patients with AD) by computing ARI (mean ± standard deviation (SD)) (Figure 2A). The most stable levels across all levels of clustering (k =2:17) were obtained for levels 2 up to 11. For this range of clustering levels (k = 2:11), there were small magnitude of the differences in case of ARI suggested that, overall, they all offered high stability. The smallest value of k where comparison between level k and k+1 was statistically significant was k = 11 (p-value < 0.001) (Figure 2A). Consequently, to define the optimum level of clustering we focused on internal validity criteria including silhouette value and Dunn index (Figure 2 (B and C)). Our results showed that clustering with three (k = 3) levels of clustering outperforms the other levels of clustering across bootstrap samples.

**Figure 2-.**
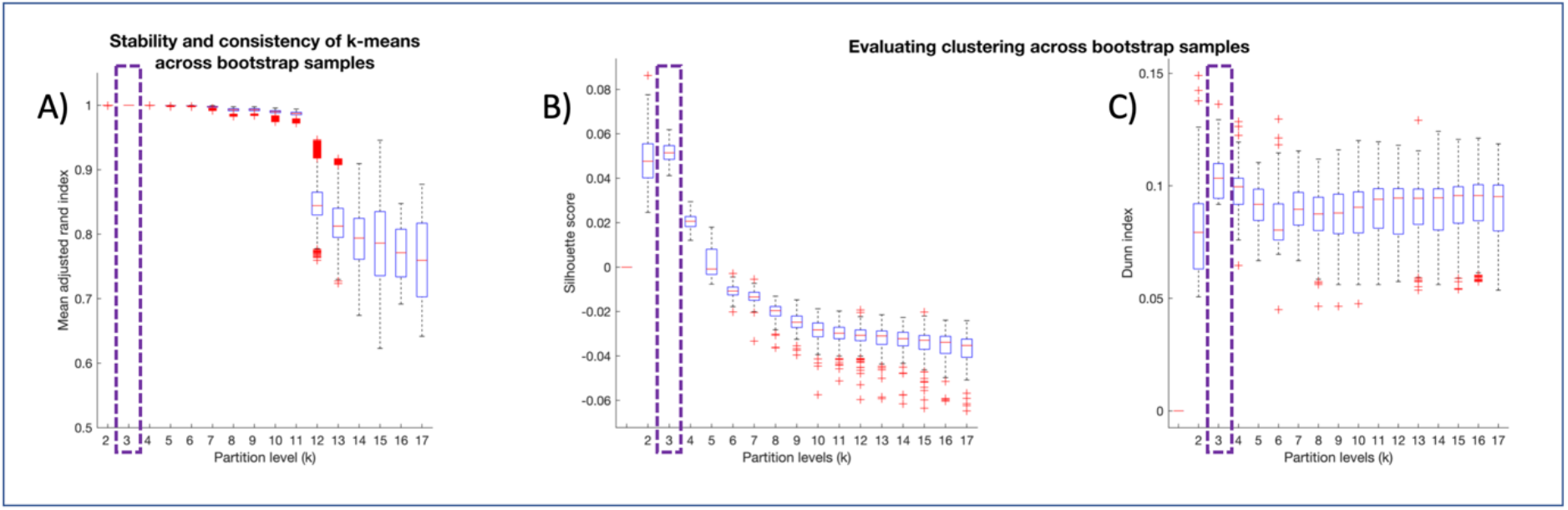
(A) Stability and consistency of all levels of clustering (k = 2:17) across bootstrap samplings was evaluated by using adjusted rand index (mean ± standard errors). The most stable levels across all clustering models (k = 2:17) were obtained for level 2 up to 11. For this range of levels of clustering (2-11), there was small magnitude of the differences in case of adjusted rand index suggested that, overall, they all offered high stability. The smallest k value where comparison between level k and k+1 was statistically significant was k = 11 (p-value < 0.001). In the following, to define the optimum level of clustering we focused on internal validity criteria including silhouette value (B) and Dunn index (C).

### 3-2- Between group differences

The optimum number of recurring FC states during rest in HC and patients with AD was determined as three (Figure 3A). Based on the clustering results, three recurring FC states which were represented by a 112× 3 matrix determined how all brain regions are contributed in each recurring FC state. Besides, we also had information related to each V1(t) (dFC pattern at time t) in each individual belonged to which recurring FC state. To identify each recurring FC state represents which brain networks or subsystems, we used Neurosynth^1^. Neurosynth is a platform for automated synthesis of fMRI data to generate probabilistic mappings between cognitive and FC states by combining meta-analysis, and machine learning techniques. First recurring FC state reflected global state of interconnections between all brain regions in line with previous studies (Cabral et al., 2017; Figueroa et al., 2019; Larabi et al., 2020). This state occurred about 46.77% of the time in HC subjects and 61.31% of the time in patients with AD (Figure 3 (A and C)). The second recurring FC state consisted of areas in FPN including the dorsolateral prefrontal cortex and posterior parietal cortex, SN such as anterior posterior cingulate cortex, and DMN including posterior cingulate cortex, precuneus, prefrontal, middle temporal and angular and supramarginal gyrus in inferior parietal lobes. We further indicated this recurring FC state as the “FPN-SN-DMN state”. FPN-SN-DMN state occurred in 32.33% of the time in HC subjects and 23.31% of the time in patients with AD (Figure 3 (A and C)). Finally, the involved brain regions in the third recurring FC state mainly located in SMN, subcortical (sub) and visual network (VIS) based on Neurosynth. This recurring FC state occurred 20.9% of the time in HC and 15.38% of the time in patients with AD (Figure 3 (A and C)). This recurring FC state was indicated further as the “SMN-sub-VIS state”. Then, we calculated the probability of occurrence, mean lifetime, and switching profiles of each recurring FC state in each individual, followed by comparing these dynamic characteristics of recurring FC states between HC and patients with AD to assess how they varied between groups.

**Figure 3-.**
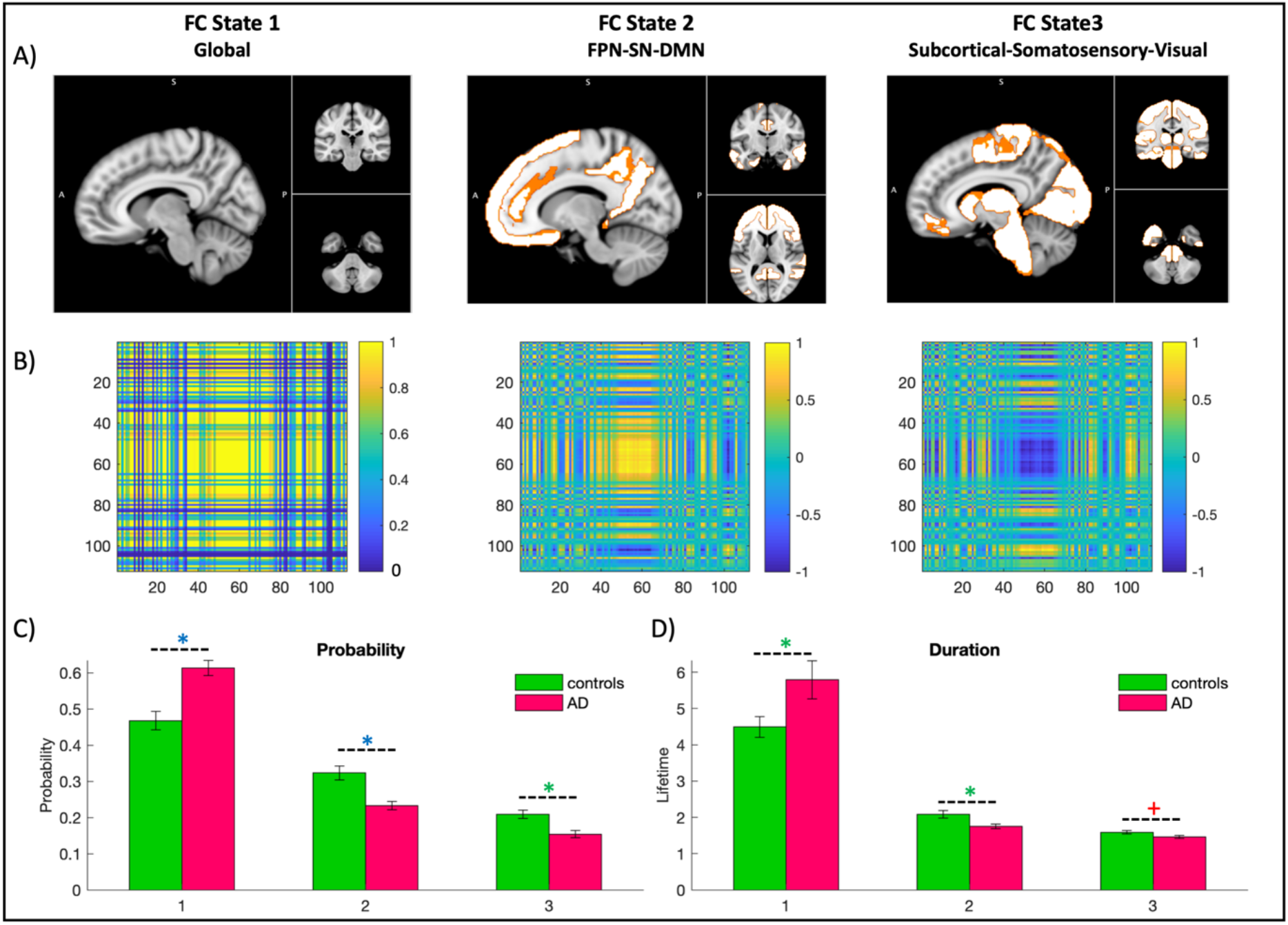
(A) Identified functional connectivity (FC) states within the most stable level of clustering, k=3 including global, FPN-SN-DMN and SMN-sub-VIS networks. All the FC state are presented in the MNI space, where functionally connected brain areas. (B) FC pattern of FC states. Results showed that in global state all brain regions were integrated and synchronized. But in the FC states 2 and 3 we observed anti-correlation pattern. For example, in FC state 2, functional interaction between DMN, FPN and SN highly correlated and these regions involved in this FC state anti-correlated with the rest of brain. Notably, involved regions in the second and third FC states were highly anti-correlated. (C) We presented the mean and standard deviation of each FC-state’s probability of occurrence and lifetime for both groups. Our analysis revealed that global FC state, which consistently appeared significantly less in HC (0.47 ± 0.026) compared to AD patients (0.61 ± 0.020), FWE corrected p-value = 1.12e^-5^. However, second and third FC states appeared statistically less in AD patients compared to HC (FPN-SN-DMN state: HC (0.32 ± 0.019), AD (0.23 ± 0.012), FWE corrected p-value = 1.43e-6; SMN-sub-VIS state: HC (0.21 ± 0.0113), AD (0.15 ± 0.010), FWE corrected p-value = 0.001). Interestingly, the most significant differences in probability of occurrence between patients with AD and HC occurred in FPN-SN-DMN state. ^+^/* Significant difference between patients with AD and controls before/after correcting for number of states 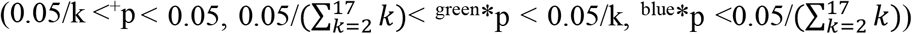. FC: functional connectivity; AD: Alzheimer’s disease; HC: healthy control; FPN: frontoparietal network; SN: salience network; DMN: default mode network; SMN: somatomotor network; sub: subcortical regions; VIS: visual network.

#### 3-2-1- probability of occurrence

We examined the probability of occurrence for each recurring FC state in each individual in both groups to explore the differences between HC subjects and patients with AD. In Figure 3C, we illustrated mean and SD of probability of occurrence for each recurring FC state in both groups. Our analysis revealed that global state consistently appeared significantly less in HC (0.47 ± 0.026) compared to patients with AD (0.61 ± 0.020), FWE corrected p-value = 1.12e^−5^. However, FPN-SN-DMN and SMN-sub-VIS states appeared statistically less in AD patients compared to HC subjects (FPN-SN-DMN state: HC (0.32 ± 0.019), AD (0.23 ± 0.012), FWE corrected p-value = 1.43e-6; SMN-sub-VIS state: HC (0.21 ± 0.0113), AD (0.15 ± 0.010), FWE corrected p-value = 0.001). Interestingly, the most significant differences in probability of occurrence between patients with AD and HC subjects with considering smallest p-value occurred in FPN-SN-DMN state.

#### 3-2-2- Lifetime

Between-group comparisons in terms of lifetime of each recurring FC state revealed that global state lasted significantly shorter in HC subjects compared to patients with AD (HC (4.49 ± 0.29), AD (5.79 ± 0.53), FWE corrected p-value = 0.036), while FPN-SN-DMN state lasted significantly longer in HC subjects compared to patient group (HC (2.09 ± 0.11), AD (1.75 ± 0.06), FWE corrected p-value = 0.014) (Figure 3D). There was no significant difference between groups in the life time of SMN-sub-VIS state (the p-value did not survive the correction for multiple comparisons (Figure 3D)).

#### 3-2-3- Switching probabilities

We also examined the transition patterns between recurring FC states by calculating the probability of being in a given FC state and switching to any of the other FC states. We illustrated the general switching pattern in patients with AD, HC subjects, and the whole group in Figures 4 (A, B and C). Here, we reported the switching probabilities that exceed a probability of 60%. On the whole-group level, the most common switching was observed from global state (63%) and SMN-sub-VIS state (66%) toward the FPN-SN-DMN state suggesting that after being in those states for some time, the areas involved in these functional networks synchronize their BOLD pattern with the FPN-SN-DMN state.

**Figure 4-.**
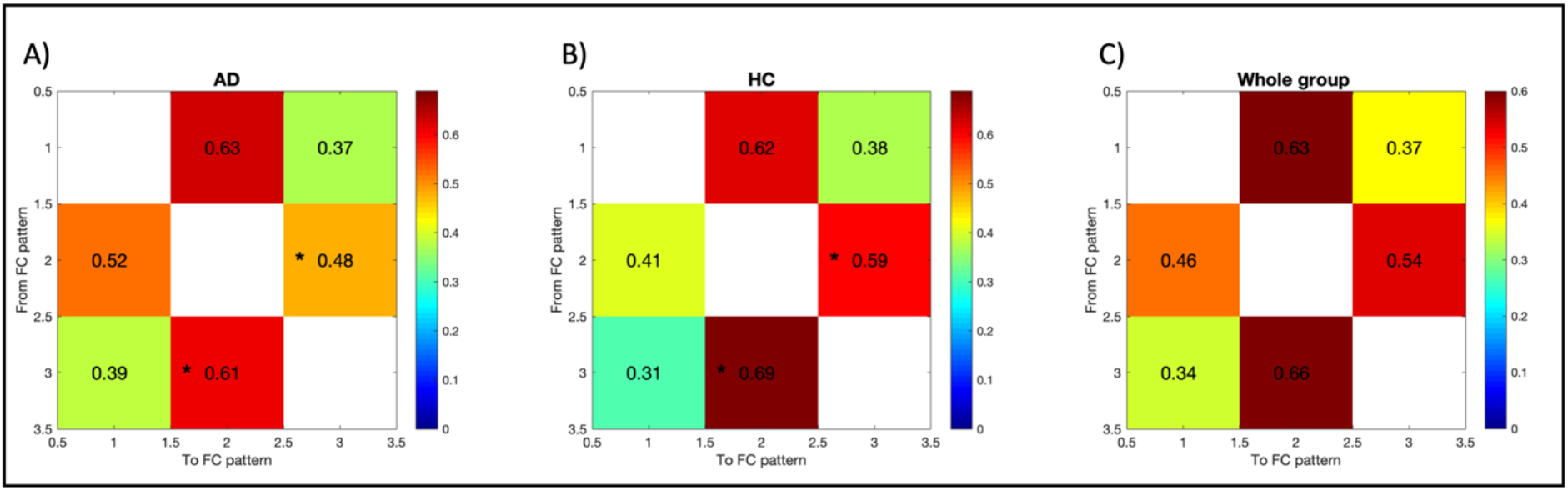
Switching probabilities for the whole group and differences between patients with AD and controls. The all switching matrices indicate the probability of, being in a given FC-state (rows), transitioning to any of the other states (columns) for AD patients (A), HC subjects (B) and the whole group (C). Significantly different transitions between both groups after correcting for multiple comparisons were identified by star. The probability of switching from FPN-SN-DMN state to SMN-sub-VIS state and vice versa were significantly less in patients with AD. Transitioning differences between groups were calculated using a permutation-based two sample t test with 10,000 permutations. FC: functional connectivity; AD: Alzheimer’s disease; FPN: frontoparietal network; SN: salience network; DMN: default mode network; SMN: somatomotor network; sub: subcortical regions; VIS: visual network.

Comparing the switching patterns between patients with AD and HC subjects, we found that, in patients with AD switching from SMN-sub-VIS state to FPN-SN-DMN state and vice versa showed a significantly lower probability compared to HC subjects (switching from FPN-SN-DMN state to SMN-sub-VIS state: AD (48%) vs. HC (59%), FWE corrected p-value = 0.0008; switching from SMN-sub-VIS state to FPN-SN-DMN state: AD (61%) vs. HC (69%), FWE corrected p-value = 0.0021). This finding suggested that in patients with AD, switching between recurring FC states appears to be disrupted. In other words, patients with AD not only experience less FPN-SN-DMN and SMN-sub-VIS states but also more frequently switched to the global state.

In addition, we also detected a higher probability of switching from SMN-sub-VIS state to FPN-SN-DMN state in both groups in comparison with switching FPN-SN-DMN state to SMN-sub-VIS state suggesting that in both groups the brain probably attempts to set FPN-SN-DMN state as the active state.

### 3-3- Replicability of the FPN-SN-DMN state

To explore the replicability of FPN-SN-DMN state as a recurring FC state with the most significant differences in probability of occurrence between groups in all levels of clustering, we first computed between-group comparisons in terms of probability of occurrence and lifetimes of each recurring FC state for each level of clustering (k = 2:17). In Figures 5 (A and B), we illustrated the k p-values obtained from group comparisons of each of k recurring FC states at k^th^ level of clustering. In each level of clustering, k hypotheses were tested which in turn increased the probability of false positive rate. We corrected the significance threshold to 0.05/k (FWE-corrected for each level of clustering) (green dashed line in Figures 5 (A and B)) and to 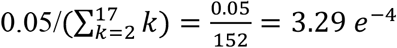 (FWE-corrected for all 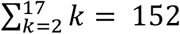 hypotheses) (blue dashed line in Figures 5 (A and B)). In Figures 5 (A and B), we used black dots for cases where there was no significant difference between groups (p-value > 0.05). The most significant results while corrected for all hypotheses in this study by using FWE rate correction (p-value < 3.29 *e*^−4^) were marked by blue stars. The significant results did not survive the correction for all hypotheses while survived the correction for each level of clustering by using FWE rate correction (3.29 *e*^−4^ < p-value < 0.05/k) were marked by green stars. Finally, the results with p-value were in range of 0.05/k < p-value < 0.05 were marked by red crosses. These p-values were less than 0.05 but not surviving the correction for level of clustering by using FWE rate corrections within each clustering level. Then, we aimed to explore among all identified recurring FC states in deferent level of clustering whether the most significant recurring FC states in each level of clustering that strongly differentiate patients with AD from HC subjects represented FPN-SN-DMN state or not? In other words, we assessed the replicability of FPN-SN-DMN state as a recurring FC state with the most significant differences in probability of occurrence between groups in all levels of clustering and its independency from what the optimum level of clustering is. Notably, we found that in the all levels of clustering (except k = 4 and 13), the most significant recurring FC state (with smallest p-value) that occurred significantly less in patients with AD compared to HC subjects consistently represented FPN-SN-DMN state (Figure 5C). In case of clustering levels of k = 4 and 13, the second most significant recurring FC state reflected FPN-SN-DMN networks. In Figure 5D, we illustrated that all mentioned significant FC sates in all levels of clustering (2:17) that occurred significantly less in patients with AD compared to HC subjects, were highly correlated with FPN-SN-DMN state identified in the most stable level of clustering (k =3) (correlation coefficient value (r) > 0.85).

### 3-4- Cognitive impairment and dynamic properties of brain recurring FC state in patients with AD

We found that patients with more cognitive decline (lower MMSE score) have significantly lower probability of occurrence of the FPN-SN-DMN state (r = 0.40, p = 0.045; Figure 6A and a significantly higher lifetime of the global state (r = -0.46, p-value = 0.021; Figure 6B). There was no further significant association between lifetime, probability of occurrence of other recurring FC states or switching probabilities between states with MMSE score.

**Figure 6-.**
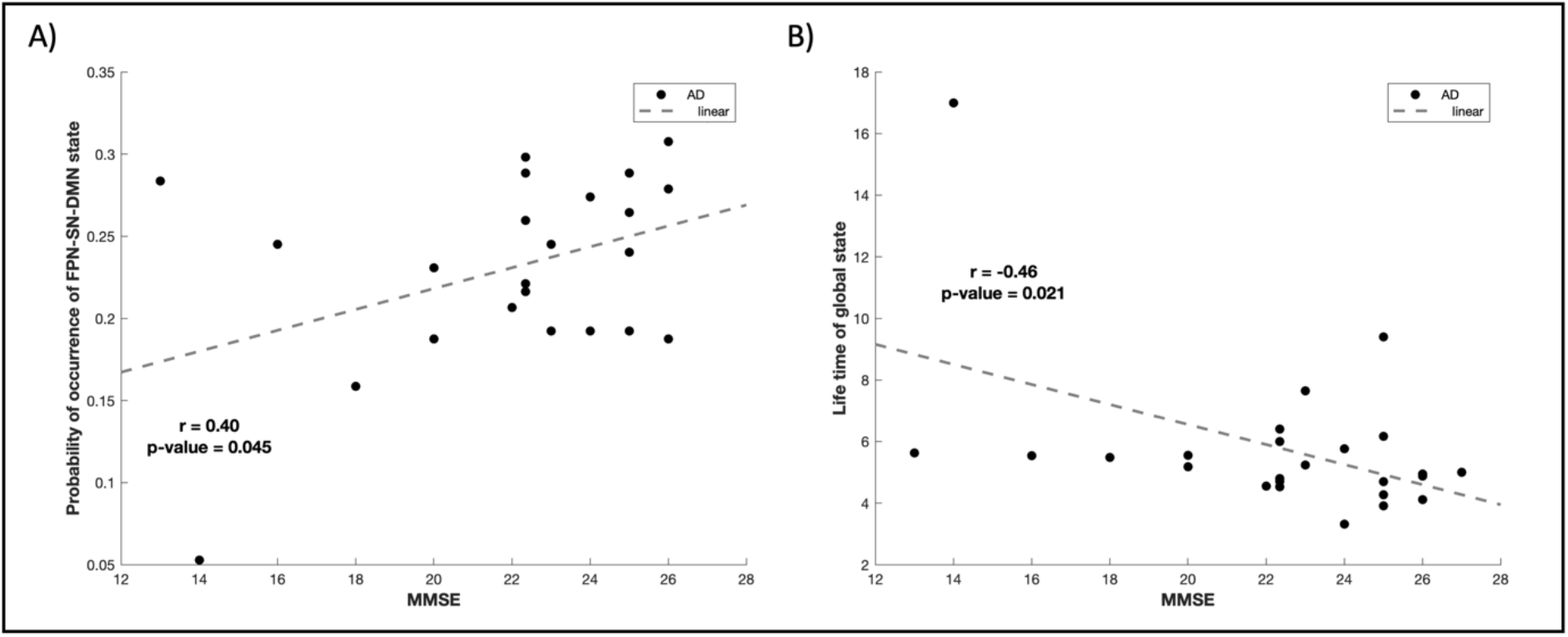
Association between cognitive impairment and dynamic properties of brain FC state in patients with AD. We found that AD patients with lower cognitive MMSE score have (A) a lower probability of occurrence of the FPN-SN-DMN state (r = 0.40, p = 0.045) and (B) a higher lifetime of the global state (r = -0.46, p-value = 0.021) than healthy control subjects. FC: functional connectivity; AD: Alzheimer’s disease; MMSE: mini-mental state examination; FPN: frontoparietal network; SN: salience network; DMN: default mode network.

## 4- Discussion

In this study, we examined how Alzheimer’s pathophysiology affects dynamic properties of resting-state brain FC. To define the optimum number of recurring FC states (k in the clustering), we first measured stability and consistency of all levels of clustering (k = 2:17) across bootstrap samples by computing ARI. Results suggested that clustering model in the range of 2 up to 11 offered high stability with small magnitude of the differences. Then, internal validity criteria including silhouette value and Dunn index were used to define the optimum level of clustering. Finally, three recurring FC states including global, FPN-SN-DMN and SMN-sub-VIS states were identified. We found that the most significant recurring FC state that strongly differentiates patients with AD from HC subjects in probability of occurrence with smallest p-value reflected FPN-SN-DMN state. FPN-SN-DMN state occurred significantly less and lasted shorter in patients with AD compared to HC subjects. This means the decreased chance in patients with AD to experience the FPN-SN-DMN state during rest. Furthermore, we revealed that identified FPN-SN-DMN state as the most significant recurring FC state in probability of occurrence between group in the optimum level of clustering can be identified in most levels of clustering as a recurring FC state with strongest difference between groups. Moreover, probability of occurrence in FPN-SN-DMN state (assessed with dFC analysis) was significantly correlated with cognitive decline (assessed by MMSE score). Consistently, AD patients had significantly lower probability of occurrence and stayed less (but not significantly) in SMN-sub-VIS state. On the other hand, patients with AD significantly showed higher probability of occurrence in the global state and spent more time in this state. We also found a negative association between cognitive decline and the lifetime of global state in patients with AD. It implies that patients with severe symptoms of AD spend more time in the global state and attempt to make brain more integrated. Our findings in impaired dynamic profile of recurring FC states in AD provide greater understanding of pathophysiology of AD.

### 4-1- Brain recurring FC states

The three recurring FC states varied in overall functional interaction patterns and dynamic properties. Except for the global state, both the FPN-SN-DMN state and the SMN-sub-VIS state contained strong anti-correlation patterns. For example, in FPN-SN-DMN state, regions involved in these three networks were highly correlated together and anti-correlated with the rest of brain. This state is characterized by positive connections between DMN and attention network and SN. This state contained FC between brain regions where AD dysconnectivity have been repeatedly reported in previous neuroimaging studies, including the posterior cingulate/precuneus (Sorg et al., 2009; Yokoi et al., 2018), anterior cingulate cortex (Amanzio et al., 2011; Brier et al., 2012), medial temporal cortex (Pasquini et al., 2019; Sorg et al., 2009), and fontal cortex (Zhao et al., 2018). Interestingly, the correlated patterns in FPN-SN-DMN and SMN-sub-VIS states were anti-correlated which indicated that AD patients significantly differ from HC subjects in probability of occurrence of these anti-correlated FC patterns. It is also noted that an important characteristic of global state is no anti-correlations. It has been suggested that global state reflects a baseline FC state in which the whole brain is going to be more integrated and segregated (Cabral et al., 2017; Figueroa et al., 2019). The global state in the present study characterized by a whole brain connectivity profile with absence of strong negative correlations that participants spend most of their time in this state. This finding is in accordance with previous findings in healthy participants (Allen et al., 2014; Cabral et al., 2017; Larabi et al., 2020), AD and dementia (Fu et al., 2019; Gu et al., 2020; Jones et al., 2012; Niu et al., 2019; Schumacher et al., 2019), aging (Tian et al., 2018; Viviano et al., 2017), and other psychiatric and neuro-degenerative diseases (Figueroa et al., 2019; Kim et al., 2017).

Furthermore, we assessed the replicability of most significant differing recurring FC state in probability of occurrence between groups, i.e FPN-SN-DMN state, across all levels of clustering. The evaluation demonstrated that in the all levels of clustering (except k = 4 and 13), the most significant recurring FC state that occurred significantly less in patients with AD compared to HC subjects, consistently represented FPN-SN-DMN networks (Figure 5C). In case of clustering levels of k = 4 and 13, the second most significant recurring FC state reflected FPN-SN-DMN state. This finding indicated that the identified recurring FC state in this study as the FPN-SN-DMN state that strongly differentiated AD patients from HC subjects were independence of how the important parameter in K means algorithm, number of clusters, were adjusted.

### 4-2- AD disrupts dynamic properties of brain recurring FC states

In our study, three recurring FC states including global, FPN-SN-DMN and SMN-sub-VIS states were identified. Among them, the most significant discriminative state was FPN-SN-DMN state in which patients with AD experience lower probability of occurrence and spend less time compared to HC subjects. Previous studies have shown that the brain regions involved in FPN, SN and DMN resting state FC networks have been affected in AD (Damoiseaux et al., 2012; Greicius et al., 2004; Sorg et al., 2009; Zhao et al., 2018). The FPN included in FPN-SN-DMN state generally also known as the central executive network and consists of the dorsolateral prefrontal cortex and posterior parietal cortex (Gong et al., 2016) activated during executive attention (Zhao et al., 2018) and working memory (Menon, 2011). Furthermore, DMN is a dominant component in FPN-SN-DMN state associated with self-related cognitive processing, such as introspection and autobiographic memory (Buckner et al., 2008), thinking about others and remembering the past and imaging the future (Andrews-Hanna, 2012). The function of DMN network appears to be crucial for mental health (Grieder, Wang, Dierks, Wahlund, & Jann, 2018). Finally, FPN-SN-DMN state included areas of the SN, a resting state FC network that facilitates switching brain activity between the FPN and DMN networks (Menon, 2011). Taken together, our results suggest that patients with AD experienced decreased ability to access brain regions involved in FPN-SN-DMN state. Additionally, we observed a significantly lower probability in patients with AD to switch from the FPN-SN-DMN state to a SMN-sub-VIS state while higher probability (not significantly) to switch to global state. The other recurring FC state that AD patients had lower probability of occurrence and spent less time compared to HC subjects are located in SMN-sub-VIS networks. Decreased ability access to brain areas were located mainly in DMN, SN, FPN, SMN, sub and VIS networks in AD patients compared to HC subjects consistent with many previous findings (Brier et al., 2012; Dai et al., 2015; Greicius et al., 2004; Gu et al., 2020; Mosimann et al., 2004; Schumacher et al., 2019). For the DMN,FPN and SN, the decreased access to these networks might be associated with impaired cognitive functions, as the DMN plays a pivotal role in essential cognitive processes (Buckner et al., 2008; Zhao et al., 2018) and especially influences memory consolidation (Greicius et al., 2004), and the FPN is involved in cognitive control and has also been reported to be also affected by AD (Zhao et al., 2018). Decreased occurrence in the SMN and VIS networks in AD patients could be related to reduced flexibility in sensory, motor, and visual functions. Interestingly, previous studies on AD and dementia with Lewy bodies (DLB) have also found recurring FC state within visual and motor networks (Schumacher et al., 2019; Sourty et al., 2016). Schumacher et al (2019) has reported a decreased ability to switch to the FC state overlapped with visual and motor networks in AD and DLB groups compared to HC subjects. In DLB patients, Sourty et al., (2016) found dynamic connectivity changes for networks related to visual processing using Hidden Markov Models. Additionally, the global state had the highest probability of occurrence in both HC and patient groups which was also reported by previous studies (Allen et al., 2014; Cabral et al., 2017; Figueroa et al., 2019; Gu et al., 2020; Larabi et al., 2020; Schumacher et al., 2019; Tian et al., 2018). Since brain global state allows for a greater range of either integration or segregation between neural networks and brain areas (Nomi et al., 2017), the high probability of occurrence of the global state may suggest that the human brain prefers switching to a state where become more integrated and organized (Nomi et al., 2017). Our results showed that the global state occurred more and lasted longer significantly in patients with AD compared to HC subjects. This finding is also consistent with previous studies that considered AD and DLB as a main cause for higher occurrence and longer lifetime in global state (Schumacher et al., 2019). However, the opposite pattern was reported in a recent study that used a similar method (sliding-window approach with a short window length (i.e., 30 s)) on AD patients (Gu et al., 2020).

Moreover, it has been suggested that global state probably reflects the global signal being combination of neural signal and artifacts such as motion, heart rate and breathing (Cabral et al., 2017; Figueroa et al., 2019; Keller et al., 2013; Larabi et al., 2020; Murphy & Fox, 2017) or a baseline FC state where all brain regions are synchronized and integrated. This finding suggested that patients with AD lost their access to FPN-SN-DMN state and switched more frequently to global brain state. We also speculated that more frequently occurrence of the global state in patients with AD might reflect a compensatory mechanism to regulate brain activity. This could be an attempt to activate FPN-SN-DMN state again after a decrease in ability to access FPN-SN-DMN state occurred in patients with AD (which might or might not be successful). However, in HC subjects, FPN-SN-DMN state switched to the SMN-sub-VIS state with higher probability than to the global state and therefore brain integration/segregation might be regulated more automatically in HC subjects which consequently lead to less need for global state.

### 4-3- Cognitive impairment and dynamic properties of brain recurring FC state in patients with AD

It is also noted that the altered dynamic properties of brain recurring FC states were correlated specifically with impaired cognitive performance in patients with AD. For example, the probability of occurrence in FPN-SN-DMN state and lifetime of the global state significantly correlated with MMSE scores (Figures 7 (A and B)). This finding demonstrated that the disruption of occurring rate or active time of a specific brain FC state were seriously correlated with the alternation of brain cognition performance in AD patients. Our findings were thus supportive of the suggestion that cognitive performance not only in healthy older adults (Cabral et al., 2017) but also in AD patients associated with dynamic profile of brain recurring FC states. In addition, since the MMSE score can reflect the severity of cognitive impairments in AD, our findings also imply that a decrease in ability to access to FPN-SN-DMN state may be closely associated with cognitive impairment symptoms, such as having trouble in remembering, learning new things, concentrating, or making decisions in AD patients. The longer life time on the global state in AD suggested that it may needs more attempts to be integrated to cope with neuropsychiatric symptoms.

### 4.4 Limitation and future work

The main strength of our study is that we focused on identification of brain recurring FC states over time at subject level instead of focusing on group representative pairwise dFC between brain regions. This gave us opportunity to compare dynamic properties of identified recurring FC states between group statistically. Although, the present work has a few limitations that should be mentioned. First, the parcellation atlas (Harvard-oxford) used to define brain regions to examine dFC may cause some anatomical bias due to different size of brain regions. In compare to functional based parcellation (Craddock, James, Holtzheimer, Hu, & Mayberg, 2012; Schaefer et al., 2018), relatively large brain regions in Harvard-oxford atlas might have less functional homogeneity within a brain region. Therefore, to avoid this potential bias, future studies can consider using other parcellations based on functional homogeneity atlas with more fine-grained brain regions. This would allow to reveal more specific recurring FC states instead of identifying a FC state that different brain resting state FC networks are involved in it. Second, since previous dFC studies on AD have found different findings and our results are both consistent and opposite with previous studies, further validation studies in large cohort of AD population are needed. Lastly, we employed the widely used tapered sliding-window approach to examine dFC in AD. Considering recent developments in this area, future studies can take into account using other developed methods to analyze dFC in AD. However, being able to discern meaningful dynamic properties in brain recurring FC states differentiating patients with AD from healthy control subjects manifests the possibilities of dFC measures to provide biomarkers for neurodegenerative disease.

### 4.5. Conclusion

This study provides new insights on aberrations in dynamic characteristic of brain recurring FC states over time in patients with AD. Our findings suggest reduced ability and flexibility of patients with AD to access FPN-SN-DMN state and switched more frequently to global brain FC state to regulate brain activity where allows for a greater range of either integration or segregation between neural networks and brain. Overall, our study demonstrated that brain recurring FC states were disrupted in AD patients, which extends the current understanding about human brain connectome in this disease, and shows the importance of evaluating dynamic variability of brain recurring FC states in AD.

## Acknowledgment

Data used in preparation of this article was obtained from the Alzheimer’s Disease Neuroimaging Initiative (ADNI) database (adni.loni.usc.edu). As such, the investigators within the ADNI contributed to the design and implementation of ADNI and/or provided data but did not participate in analysis or writing of this report. A complete listing of ADNI investigators can be found at: http://adni.loni.usc.edu/wpcontent/uploads/how_to_apply/ADNI_Acknowledgement_List.pdf. Clinical data collection and sharing for this project was funded by the Alzheimer’s Disease Neuroimaging Initiative (ADNI) (National Institutes of Health Grant U01 AG024904) and DOD ADNI (Department of Defense award number W81XWH-12-2-0012). ADNI is funded by the National Institute on Aging, the National Institute of Biomedical Imaging and Bioengineering, and through generous contributions from the following: AbbVie, Alzheimer’s Association; Alzheimer’s Drug Discovery Foundation; Araclon Biotech; BioClinica, Inc; Biogen; Bristol-Myers Squibb Company; CereSpir, Inc; Cogstate; Eisai Inc; Elan Pharmaceuticals, Inc; Eli Lilly and Company; EuroImmun; F Hoffmann-La Roche Ltd and its affiliated company Genentech, Inc; Fujirebio; GE Healthcare; IXICO Ltd.; Janssen Alzheimer Immunotherapy Research and Development, LLC; Johnson and Johnson Pharmaceutical Research and Development LLC; Lumosity; Lundbeck; Merck and Co, Inc; Meso Scale Diagnostics, LLC; NeuroRx Research; Neurotrack Technologies; Novartis Pharmaceuticals Corporation; Pfizer Inc; Piramal Imaging; Servier; Takeda Pharmaceutical Company; and Transition Therapeutics. The Canadian Institutes of Health Research is providing funds to support ADNI clinical sites in Canada. Private sector contributions are facilitated by the Foundation for the National Institutes of Health (www.fnih.org). The grantee organization is the Northern California Institute for Research and Education, and the study is coordinated by the Alzheimer’s Therapeutic Research Institute at the University of Southern California. ADNI data are disseminated by the Laboratory for Neuro Imaging at the University of Southern California.

## Footnotes

1 http://neurosynth.org/decode/

